# The enzymatic and neurochemical outcomes of a mutation in Mexican cavefish MAO reveal teleost-specific aspects of brain monoamine homeostasis

**DOI:** 10.1101/2022.10.10.511577

**Authors:** Constance Pierre, Jacques Callebert, Jean-Marie Launay, Sylvie Rétaux

## Abstract

Monoamine oxidases (MAO; MAO-A and MAO-B in mammals) are enzymes catalyzing the degradation of biogenic amines, including monoamine neurotransmitters. In humans, coding mutations in MAOs are extremely rare and deleterious. Here, we assessed the structural and biochemical consequences of a point mutation (*P106L*) in the single *mao* gene of the blind cavefish *Astyanax mexicanus*. This mutation decreased mao enzymatic activity by ~3-fold, probably as a result of decreased flexibility in one of the three loops forming the entrance of the active site, thus reducing the access of substrates. HPLC measurements in brains of mutant and non-mutant larvae and adults of the cave and surface morphs of the species showed major disturbances in serotonin, dopamine and noradrenalin (and metabolites) contents in mutants, demonstrating that the P106L *mao* mutation is fully responsible for monoaminergic disequilibrium in the P106L *mao* mutant cavefish brain. The outcomes of the mutation were different in the posterior brain (containing the raphe nucleus) and the anterior brain (containing fish-specific hypothalamic serotonergic clusters), revealing contrasting properties in neurotransmitter homeostasis in these different neuronal groups. We also discovered that the effects of the mutation were partially compensated by a decrease in activity of the tph, the serotonin biosynthesis rate-limiting enzyme. Finally, the neurochemical outcomes of the *mao* P106L mutation differed in many respects from a treatment with deprenyl, an irreversible MAO inhibitor, showing that genetic and pharmacological interference with MAO function are not the same. Our results shade light on our understanding of cavefish evolution, on the specificities of fish monoaminergic systems, and on MAO-dependent homeostasis of brain neurochemistry in general.

## Introduction

Monoamine oxidase (MAO) is a flavoprotein that catalyzes the oxidative deamination of biogenic amines – monoamine neurotransmitters and dietary amines. It is located at the outer membrane of mitochondria in neuronal, glial and other cells, in platelets, liver and kidney for example (Berry et al., 1994; Levitt et al., 1982). In mammals, MAO exists under two isoforms that share 70% identity in primary sequence and differ in terms of substrate and inhibitor specificities (Bach et al., 1988). MAO-A prefers serotonin (5-hydroxytryptamine, 5-HT) and noradrenalin (NA) as substrates and is sensitive to the irreversible inhibitor clorgyline, while MAO-B prefers phenylethylamine and benzylamine as substrates and is more sensitive to the selective inhibitor deprenyl. Dopamine (DA) and tyramine are common substrates for both isoenzymes (Shih et al., 1999). The two *MAO* genes are closely linked on chromosome X and share the same exon-intron organization, suggesting that they have evolved from an ancestral gene by tandem duplication (Grimsby et al., 1991).

The enzyme’s biomedical relevance is well supported by the proven efficacy of pharmacological inhibitors in treating e.g. depression (Youdim and Bakhle, 2006) and the major neurological defects caused by its malfunction or de-regulation. In humans, only two families were reported with mutations in *MAO-A*, and they show cognitive and mental retardation, with severe behavioral disturbances including violent behavior and impulsive aggressiveness (Brunner et al., 1993; Piton et al., 2013). In mice, *MAO-A* and *MAO-B* knock-out lines with elevated brain monoamine levels also display complex behavioral alterations similar to the human “Brunner syndrome”, and modifications of stress response and anxiety-like behavior, respectively (Cases et al., 1995; Grimsby et al., 1997). The double *MAO-A/B* knock-out mice show hallmarks of autism spectrum disorder and greater neuropathological alterations than single KO mice (Bortolato and Shih, 2011). Moreover, altered *MAO* gene promoter methylation and its interaction with environmental factors emerges as an important mechanism in the etiology of several mental disorders (Ziegler and Domschke). Finally, recent work suggests that besides its neurophysiological aspects, MAO also plays important roles in tumorigenesis, diabetes, obesity and cardiovascular disease (Deshwal et al., 2018; Shih, 2018).

The first fish *mao* was cloned from trout and displays about 70% identity with MAO-A and MAO-B (Chen et al., 1994). In fact, teleost fish have only one *mao* in their genomes. Fish *mao* is homologous to the mammalian *MAOs*, is expressed in various tissues like in mammals, and degrades tyramine, serotonin, phenylethylamine (PEA) and dopamine (Aldeco et al., 2011; Anichtchik et al., 2006; Chen et al., 1994; Setini et al., 2005). On the other hand, some authors have reported that DA is a poor substrate for zebrafish mao (Sallinen et al., 2009). The teleost mao seems to exhibit functional properties that are more similar to those of mammalian MAO-A (Aldeco et al., 2011; Sallinen et al., 2009), and zebrafish *mao* knock-out line shows various neurological alterations (Baronio et al., 2021).

In the blind and depigmented, cave-dwelling form of the fish species *A. mexicanus* (Elliott, 2018; Mitchell et al., 1977), a natural point mutation in the *mao* gene (P106L) has been reported (Elipot et al., 2014). The substituted proline 106 is homologous to the proline 114 and 105 in human MAO-A and -B, respectively. Therefore, through comparative studies with its river-dwelling conspecific, the cavefish natural *mao* mutant could help us better understand the properties of fish MAO and the impact of MAO deficiencies in the vertebrate brain. Moreover, as the zebrafish *mao* knock-out is non-viable (Baronio et al., 2021), the *Astyanax mao* P106L is an excellent tool to study the effect of MAO deficiency in fish. The consequences of the P106L mutation in cavefish *mao* on relevant physiological, behavioral and developmental traits have been studied previously (Pierre et al., 2020). Surprisingly, *mao* P106L has little deleterious phenotypic effects and mainly alters stress response in cavefish as compared to surface fish. In the present study, we focused on the impacts of the P106L *mao* mutation on enzymatic activity and brain neurochemistry. We report that P106L leads to a 3-fold reduction of enzymatic activity, thus corresponding to a partial loss-of-function mutation, which can be explained by *in silico* structure-function analyses. In the cavefish brain, P106L causes a major disequilibrium in 5-HT, DA and NA neurotransmission indexes, but we found that 5-HT homeostasis is partially compensated for at synthesis level. Finally, we showed that pharmacological treatments reducing MAO activity have markedly different outcomes than the P106L genetic mutation in cavefish, suggesting that plastic and compensatory mechanisms must occur and may buffer the deleterious effects of mutation.

## Methods

### Fish husbandry

Laboratory stocks of *A*. *mexicanus* surface fish and cavefish (Pachón population) were obtained in 2004–2006 from the Jeffery laboratory at the University of Maryland, College Park, MD, USA and were since then bred in our local Gif facility. Fish were maintained at 23–26 °C on 12:12-h light: dark cycle and they were fed twice a day with dry food. The fry were raised in Petri dishes and fed with micro-worms after opening of the mouth (~6dpf, days post-fertilization). Animals were treated according to the French and European regulations for handling of animals in research. SR’s authorization for use of *Astyanax mexicanus* in research is 91-116 and the Paris Centre-Sud Ethic Committee protocol authorization number related to this work is 2017-05#8604. The animal facility of the Institute received authorization 91272105 from the Veterinary Services of Essonne, France, in 2015.

### Fish lines

To obtain Pachón CF without the P106L mutation, we crossed heterozygotes fish identified among our lab Pachón breeding colony, taking advantage that the P106L mutation is not fixed in the Pachón population (see (Pierre et al., 2020)). To obtain a SF line carrying the mutation, a cross between a SF (wild type *mao*) and a CF carrying the P106L (homozygote mutant) was followed by 4 backcrosses with SF (see (Pierre et al., 2020). Then, to obtain SF homozygote mutants, we intercrossed the last generation together. Note that the generation time between the spawn of the n generation and the spawn of the n+1 generation was about 8 months.

### *mao* P106L allele genotyping

To genotype adults, we took fin-clips and performed a lysis with proteinase K in lysis buffer (100mM Tris; 2mM EDTA; 0.2% Triton; 0.01μg/μl PK), followed by a PCR (primer F-GGGAAATCATATCCATTCAAGGGG; primer R-CTCCATGCCCATCTTGTCCATAG), and a purification of DNA (NucleoSpin® Gel and PCR Clean-up). We used the genotyping service of Eurofins Genomics. Homozygotes and heterozygotes at position 106 were easily detected and identified on sequence chromatograms. We did not genotype larvae as they were obtained from genotyped groups of adults breeding fish.

### Sampling for biochemistry

6dpf larvae were anesthetized in water at 1°C, and the head was cut from the body. A sample (n=1) was formed with 15 heads or bodies in 400μl of HCl (10^−3^M). The same protocol was used for 1month-old larvae, but samples were made up of 10 heads or bodies. Adults (5months-old) were quickly anesthetized in water at 1°C. Then the head was cut with a scalpel, the brain dissected out and cut in two halves to separate the anterior part (with hypothalamus serotonergic clusters) and the posterior part (with raphe nuclei). The 2 parts of the brain were individually placed in 400μl of HCl (10^−3^M).

### HPLC and enzymatic activities

For HPLC: before analysis, tissues were crushed and centrifuged at 20.000g for 1h. The supernatant was analyzed by fluorimetry for serotonin (HPLC), coulometry for catecholamines (HPLC).

For enzymatic activities: MAO activity was measured as described in (Elipot et al., 2014), using tyramine as substrate. TPH activity was measured by radio-enzymatic assay. Tissue homogenate (25 μL) was added to a reaction mixture containing 0.05 mM tryptophan, 50 mM Hepes (pH 7.60), 5 mM DTT, 0.01 mM Fe(NH4)2(SO4)2, 0.5 mM 6-MPH4, 0.1 mg/ml catalase, and 3H-tryptophan (1 μCi/reaction). The enzymatic reaction was allowed to proceed at 37°C for 20 min. Unreacted tryptophan and the product 5-HTP were adsorbed with 500 μl of 7.5% charcoal in 1 M HCl at the end of the incubation. The samples were thoroughly vortexed and centrifuged at 14,000 × g for 2 min. The supernatant (350 μL) was centrifuged, and a 200-μl aliquot of the final supernatant was added to 3 mL of scintillation fluid and the radioactivity was measured by a liquid scintillation counter (Beckman LS 6000SC). Blank values were obtained by performing the reaction in the absence of tissue homogenate and in the presence of boiled homogenate. The counts per minute were converted to pmoles of 5-HTP formed per milligram of tissue per hour using the formula described by (Vrana et al., 1993).

### Deprenyl treatments

Adults were placed in water containing 10μM of Deprenyl for 5hrs for an acute treatment.

### 3D Modeling

3D Modeling was performed with iTasser (https://zhanglab.ccmb.med.umich.edu/I-TASSER/), 3D alignment with TM-align, and secondary structure predictions and normalized B-score with ResQ (Yang et al., 2016; Yang et al., 2015).

### Statistical analyses

No statistical method was used to predetermine sample size. No data were excluded from analysis, and sample allocation was random after genotyping. Sex was not considered in the analyses as it impossible to determine sex in *Astyanax mexicanus* before the age of 6-7 months without dissection/sacrifice. Analyses were not be blinded (note that for anatomical analyses, brains from SF or CF are easily recognizable by eye size and presence of pigmentation). Statistical analyses were performed with BiostaTGV (https://biostatgv.sentiweb.fr/), using non-parametric Mann-Whitney tests (normal distribution was not tested). All graphs show mean±sem, and the number of samples included is systematically indicated on graph bars. P values are shown. In all figures, *** p<0.0001, ** p<0.001 and * p<0.01.

## Results

### The P106L mutation severely decreases the enzymatic activity of *A. mexicanus* MAO

A previous study had measured MAO enzymatic activities on *A. mexicanus* surface fish (SF) and cavefish (CF) originating from the Pachón cave (Elipot et al., 2014). However, we have recently discovered that the P106L mutation is not fixed in the Pachón CF population (Pierre et al., 2020). Therefore, the Pachón CF used in the previous study were probably a mix of mutants, non-mutants and heterozygotes, prompting the need to assess MAO enzymatic activity on samples with homogeneous genetic backgrounds. Here, we measured MAO enzymatic activities on 6dpf brain extracts from P106L homozygous mutants, heterozygous and non-mutant CF larvae obtained from crosses. The P106L mutation caused a ~3-fold reduction of MAO enzymatic activity, using tyramine as a substrate (Fig 1A). Consistently, this reduction was stronger than the 2-fold reduction we previously reported on samples that were probably non-homogeneous genetically. Of note, MAO activity in brains from heterozygotes was also slightly reduced as compared to controls (15.1±0.39, n=3; p=0.057), probably in line with the fact that MAO functions as a dimer.

**Figure 1.**
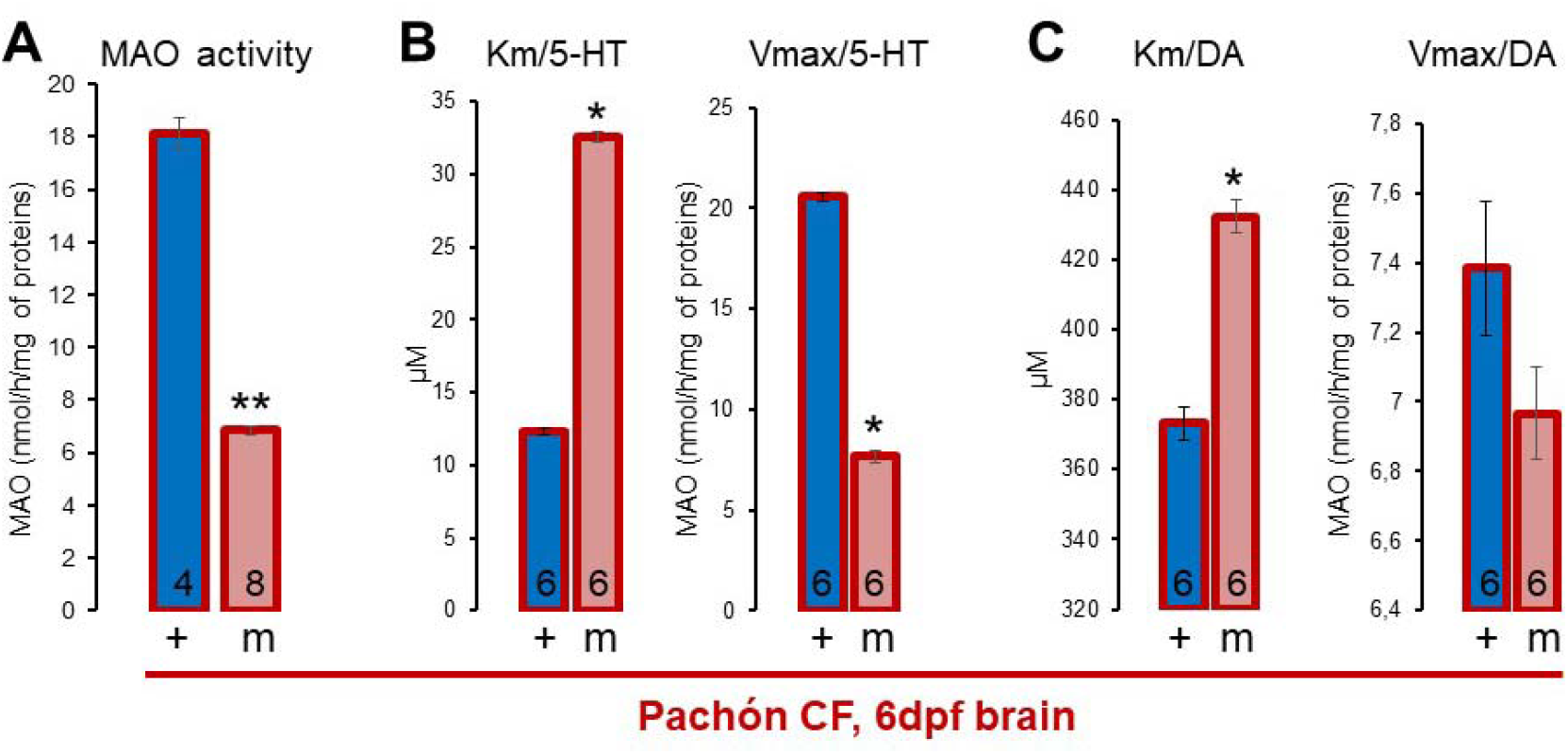
Enzymatic properties of P106L mutant and non-mutant MAO. (A) Measures of enzymatic activities in brains of 6dpf cavefish that are homozygotes *P106L MAO* mutants (m) or non-mutants (+). (B, C) Measures of K_m_ and V_max_ on serotonin (B) or dopamine (C) as a substrate. In this and subsequent figures, all graphs show mean±sem, the number of samples included is indicated on bars, and p values are shown (*** p<0.0001, ** p<0.001 and * p<0.01; Mann Whitney tests). The outlines of the histogram bars indicate the morphotype (SF, blue and CF, red) and the colors inside bars indicate the genotype (+/wildtype in blue; *mao* P106L mutant in red).

Further, we determined the Km and the Vmax of the enzyme its serotonin substrate in brain extracts from P106L *mao* mutant and wildtype *mao* adult cavefish (Fig 1B). In mutant extracts the Km was 2 to 3-fold higher, indicating a decreased affinity for the substrate, and the Vmax was 2 to 3-fold lower, in concordance with the decreased MAO activity described above. We also determined the Km and the Vmax of the enzyme for dopamine (Fig 1C). As expected, the affinity of MAO was much lower (40 times) than for serotonin (Sallinen et al., 2009). The effect of the P106L mutation was also milder on the affinity for dopamine (−13% in mutant brains), and the mutation did not significantly affect the Vmax.

### The P106L mutation probably limits entrance of the substrate to the enzyme active site

To get insights on how the *mao* P106L mutation affects the structure-function of the enzyme, we accessed 3D-modeling of the mutant and non-mutant proteins. First, a 3D model of *A. Mexicanus* MAO was obtained with iTasser (Fig 2A). Even if the P106L mutation should theoretically remove a bend in the polypeptide chain, the protein modeled with either a proline or a leucine in position 106 had the same Cα trace and perfectly overlapped, with a TM score between 0.995 and 1 depending on the models used (Fig 2B). The TM score is comprised between 0 and 1, with the value 1 indicating a perfect match between two structures. Thus, the P106L mutation did not change the structure of the MAO protein.

**Figure 2.**
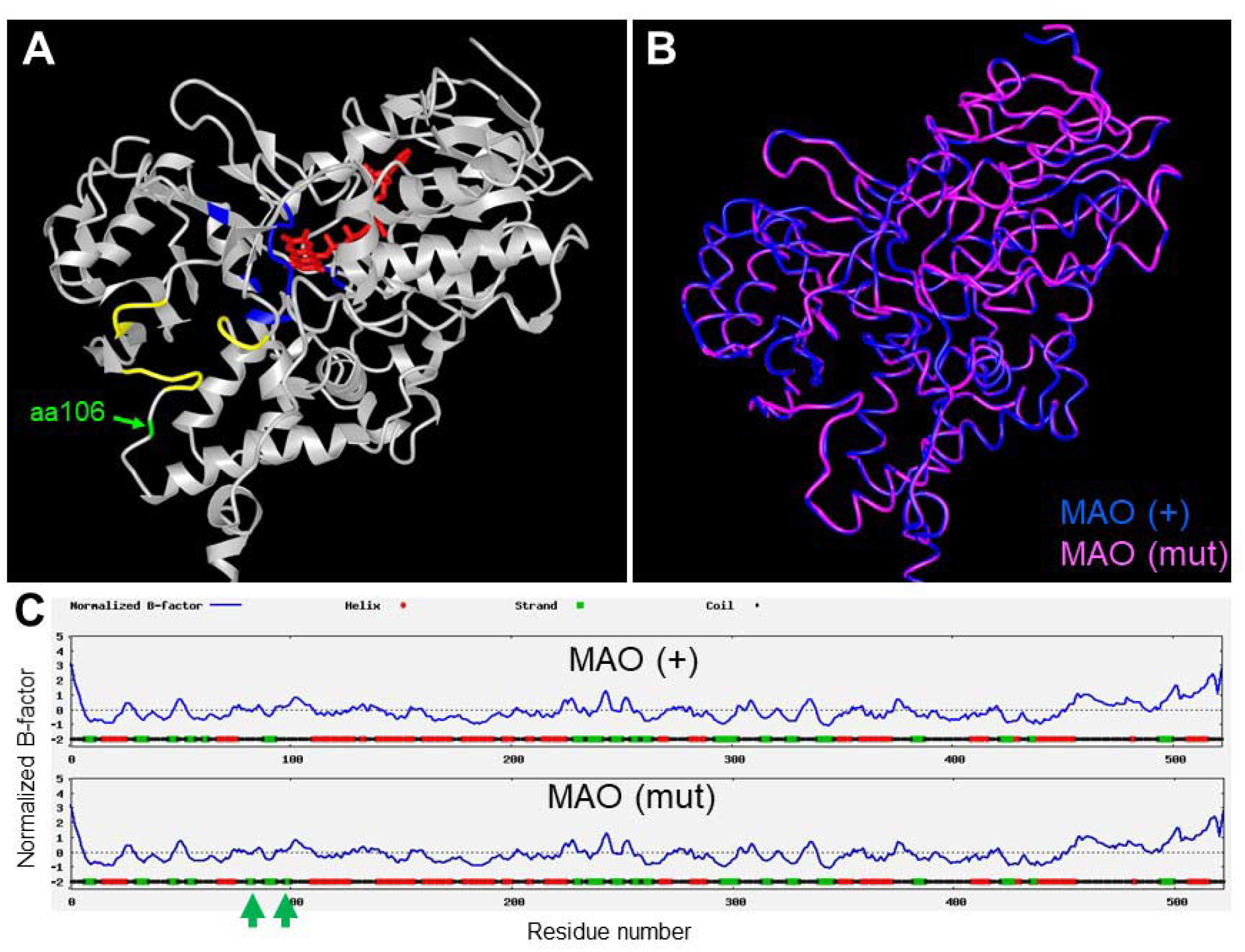
Structure of P106L mutant and non-mutant MAO proteins. (A) 3D model of the globular part of wild-type *Astyanax mexicanus* MAO, with FAD cofactor in red. Blue amino acids belongs to the active site and bind serotonin (Binda et al., 2002; Veselovsky et al., 2004), the three loops in yellow form the entrance to the active site (Son et al. 2008). The position 106 is in green (arrow). (B) P106L mutant (pink) and non-mutant MAO (blue) structure superposition. (C) Normalized B-factor (blue line) and secondary structure prediction along the protein sequence for the two MAO proteins. The normalized B-score is calculated with local variations of modeling simulations. If the normalized B-score is under zero, the residue is defined as stable, and flexible otherwise.

In human MAOs, the entrance channel for the substrate towards the deep active site is formed by 3 loops (V93-E95; Y109-P112; F208-N212) (Son et al., 2008) (see Fig 2A). Since this entrance is narrow, conformational fluctuations of the 3 loops are necessary to let the substrates enter the active site (Binda et al., 2002; Son et al., 2008). Mutations which cause a rigidification of the loop have been associated with a reduction of the enzymatic activity (Son et al., 2008).

*A. mexicanus* MAO showed a structure very similar to human MAOs, with TM scores of 0.97415 and 0.94971 when compared to MAO-A and MAO-B, respectively. The proline106 in *Astyanax* MAO is homologous to the proline114 in human MAO-A, so that the P106L mutation found in cavefish is located at the edge of one of the 3 loops (Fig 2A). In terms of secondary structure, the predictions obtained with ResQ showed two additional strands formed next to the position 106 in the mutant protein (Fig 2C). Helix and strands are more rigid than unfolded coil. Thus, combining our analyses in *Astyanax* MAO together with previous studies on human MAO-A (Son et al., 2008) suggested that the P106L mutation could rigidify one of the loops forming the entrance of the active site, and therefore limit the access of substrates. It should be noted, however, that the normalized B-score whose calculation is based on local variations of modeling simulations, and so is an indicator of the flexibility, is not very different between the two proteins (Fig 2C).

In sum, both biochemical evidence (low affinity for substrate, Fig 1B) and structural evidence (rigidity of substrate entrance loop, Fig 2) suggest that the mechanism leading to reduced enzymatic activity of MAO P106L is a limitation of the substrate’s accessibility to the active site.

Finally, to get insights on the severity of the P106L mutation from another, independent, perspective, we assessed its pathogenicity score in MutPred2 (Pejaver et al., 2017). MutPred2 is a machine learning-based predictor of the impact of missense variants in humans. The cavefish *mao* P106L mutation had a MutPred2 score, i.e., the probability that the amino acid change is pathogenic, of 0.456. This is close to the pathogenicity threshold (0.5). Of note, we have incidentally found another non-synonymous mutation just next to position 106, M107I, in a wild-caught surface fish individual (Pierre et al., 2020). By comparison, this mutation had a MutPred2 score of 0.247, suggesting that contrarily to P106L, the M107I variant found in a surface fish is not deleterious for the enzyme function.

Altogether, these data provide structural, biochemical and computational evidences that the cavefish *mao* P106L mutation corresponds to a partial loss-of-function mutation. Below, we further analyzed its consequences on cavefish brain neurochemistry.

### The P106L mutation affects monoamines homeostasis

#### The P106L mutation increases monoamine levels and decreases metabolite levels in the brain, in a region-specific manner

To assess the consequences of decreased MAO enzymatic activity on brain neurochemistry, we measured monoamines and metabolites levels in heads and bodies of 6dpf larvae and brains of 5mpf (=young adults) *A. mexicanus*. We reasoned that comparing surface fish and cavefish, either with wildtype, non-mutant *mao* or with the P106L mutation would enable us to disentangle the effect of the morphotype (surface versus cave) and the effect of the mutation (*mao* wildtype versus P106L) on brain monoamines.

As expected, in larval heads the levels of the 3 monoamines 5-HT (serotonin), NA (noradrenalin) and DA (dopamine) were higher in P106L *mao* mutants than in non-mutant samples (Fig 3A). The P106L mutation increased by 1.26, 1.15 and 1.30 times the levels of 5-HT, NA and DA, respectively. Conversely, the levels of 5-HT and DA metabolites (5HIAA, DOPAC and HVA) were lower in CF *mao* mutants. The NA metabolite VMA was non-detectable. Importantly, the levels of 5-HT and NA were identical in SF and non-mutant CF. The slight difference in DA content between SF and non-mutant CF observed in 6dpf larvae disappeared at older stages (1 month-old and 5 month-old) (Fig 3C and Fig 1 Supp).

**Figure 3.**
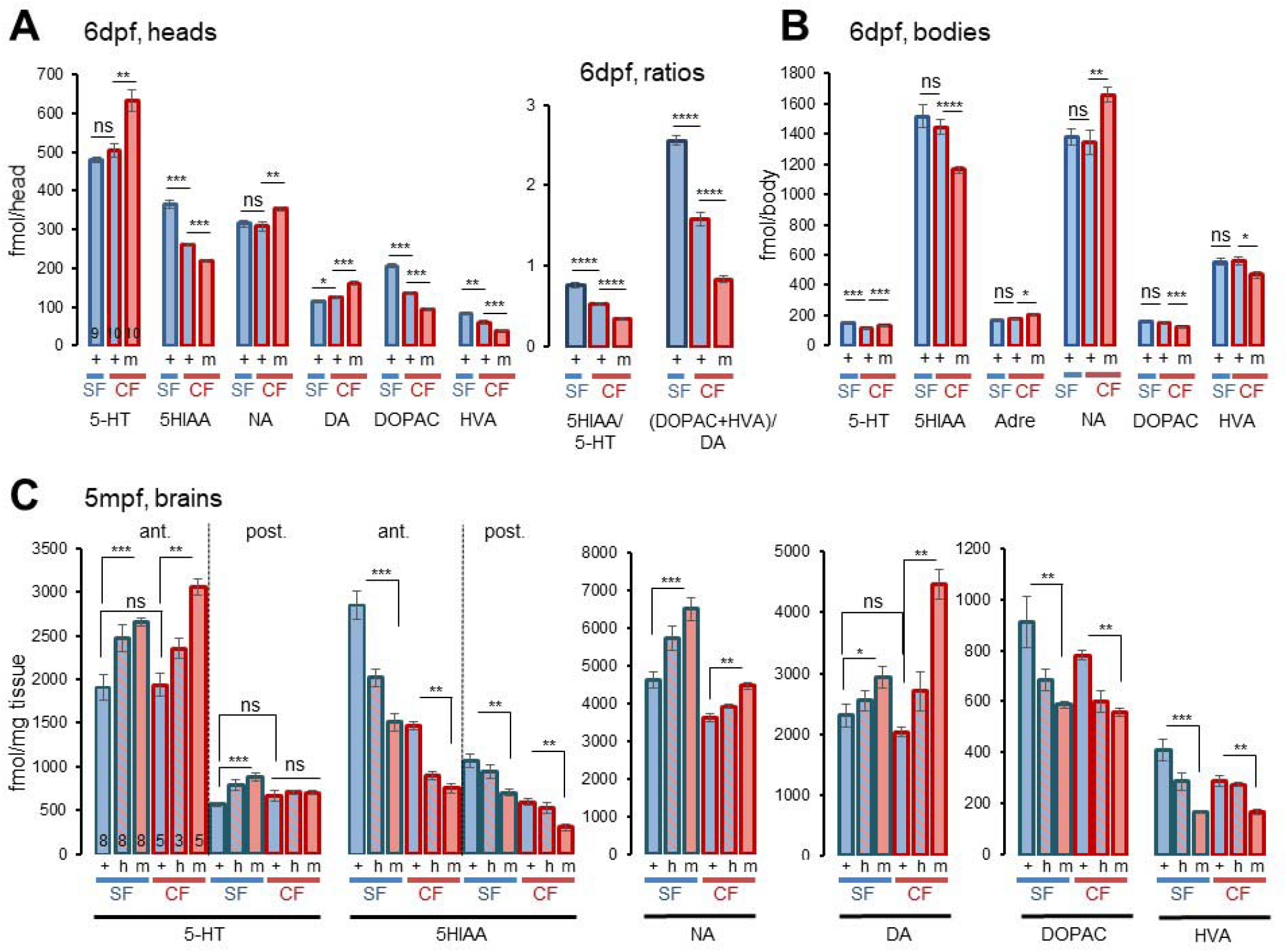
Monoamine and metabolite levels in *mao* P106L mutant or non-mutant *A. mexicanus*, surface fish and Pachón cavefish. On all graphs, the mutant (m), non-mutant (+) or heterozygous (h) state of the *mao* P106l mutation is indicated, as well as the morph (SF or CF), the type of sample (larval brain, body, or anterior/posterior adult brain) and the molecule measured. (A) Measures of monoamines and metabolites in heads of 6dpf larvae and calculated ratios 5-HIAA/5-HT and (DOPAC+HVA)/DA in SF and CF, non-mutant (+) or homozygote *mao* P106L mutants (m). Each sample (n=1) corresponds to 15 mixed heads of the same genotype. (B) Measures of monoamines and metabolites in bodies of 6dpf larvae in SF and CF, non-mutant (+) or homozygote *mao* P106L mutants (m). Each sample (n=1) corresponds to 15 mixed bodies of the same genotype. (C) Measures of monoamines and metabolites in the brain of 5 month-old fishes in SF and CF, non-mutant (+), homozygote *mao* P106L mutants (m) and heterozygotes (h).

The index of neurotransmission is often estimated by the metabolite/neurotransmitter ratio (inversely proportional to the “strength” of neurotransmission or signaling). In the heads of 6dpf larvae, the 5HIAA/5-HT and (DOPAC+HVA)/DA ratios were lower in CF, both mutant and non-mutant, and were further decreased by the mutation within CF (Fig 3A, right). This suggested a very low 5-HT and DA index and thus a strong 5-HT and DA neurotransmission in P106L *mao* mutant CF larval heads.

We also measured monoamine and metabolites in bodies of 6dpf larvae. There, the P106L mutation also increased by 1.17, 1.23 and 1.17 times the levels of 5-HT, NA and Adrenaline, respectively (Fig 3B). The metabolites 5HIAA, DOPAC and HVA were increased. VMA and DA were non-detectable in bodies. The only morph-dependent difference between non-mutant CF and SF was a higher level of 5-HT in SF. Again, this difference between non-mutant CF and SF disappeared in 1 month-old fish (Fig 3C and Fig 1 Supp).

The fish brain serotonergic system is composed of two groups of clusters of neurons, in the hypothalamus and in the raphe, respectively. Hypothalamic and raphe clusters are distinct in terms of gene expression and transcriptional factors involved during their development (Lillesaar, 2011). We therefore hypothesized that the P106L *mao* mutation could have distinct consequences in the two groups of serotonergic clusters and we analyzed separately the anterior (including the hypothalamic clusters) and the posterior (including the raphe nuclei) parts of the brain in 5 months-old fish (a cut at the midbrain/hindbrain junction was possible on older, 5mpf brains). At this age, for the two brain parts, serotonin levels were identical in SF (wildtype) and non-mutant CF (Fig 3C). This demonstrated that all the variations described below between mutant *mao* CF and wildtype *mao* SF/CF are fully and exclusively due to the P106L mutation.

Regarding 5-HT, in cavefish carrying the P106L mutation serotonin levels were increased by 1.58 times in the anterior brain but were unaffected in the posterior brain. In surface fish however, the P106L mutation led to an increase in 5-HT levels in both the hypothalamus and the raphe, thus uncovering a morph-specific difference in serotonin metabolism in the raphe. In both morphs, the heterozygotes showed intermediate levels between homozygote mutants and wildtype individuals. On the other hand, P106L decreased 5HIAA levels in both parts of the brain and in both morphs. This was unexpected but consistent with previous studies (Elipot et al., 2014) where the difference in 5-HT levels reported between SF and CF (non-genotyped) were only different in the anterior part of the brain, whereas the 5HIAA levels were different in the two parts of the brain.

Regarding DA and NA, we examined 5mpf whole brain contents. The two morphs showed very similar variations according to *mao* genotypes. In both SF and CF, the P106L mutation increased NA and DA levels, and decreased the DA metabolites (Fig 3C). Like for serotonin, heterozygotes showed intermediate levels between homozygote mutants and wildtype individuals.

Altogether, these data suggest that the P106L *mao* mutation causes major modifications in the brain neurochemistry of cavefish. The subtle but significant differences we observed between anterior and posterior parts of the brain also suggest that 5-HT hypothalamic and raphe neurons “react” differently to compromised MAO activity. Further, the differences observed between cavefish and surface fish carrying the P106L *mao* mutation suggest that the morph-specific genetic background has some influence on monoamine metabolism.

Finally, vertebrates including teleosts show daily variations of monoamine levels (Fingerman, 1976; Kahn and Joy, 1988). Yet *Astyanax* cavefish show a disruption of their endogenous biological clock and its entrainment by light (Beale et al., 2013; Mack et al., 2021; Moran et al., 2014) and *mao* P106L alters brain monoamine levels (above), prompting us to investigate whether circadian variation in monoamine levels would persist in mutants. Fish aged 5mpf were entrained during one week on a 12h:12h light-dark (LD) regime, and monoamine levels were assayed every 2h over a 26h period. As expected, cyclic variations of monoamines and metabolites were present in *Astyanax* surface fish and non-mutant cavefish, suggesting that daily variations are unaltered in the cave morph under these conditions of LD entrainment (Suppl Fig 2 and 3). However and strikingly, rhythmicity was abolished in cavefish carrying the *mao* P106L mutation. These data suggest that in cavefish, daily variations in monoamines can still be entrained by light despite the defective circadian clock, and that this process heavily relies on MAO-dependent regulatory mechanisms.

#### P106L mutant CF show a reduced TPH activity

The enzymatic activity of MAO was reduced 2.64 times by the P106L mutation in cavefish (Fig 1A). However, the levels of neurotransmitters were more modestly increased (1.58X for 5-HT in the anterior brain; 2.20X for DA)(Fig 3). We therefore wondered if a compensation could occur at the level of neurotransmitters synthesis. We measured and compared the activity of TPH (Tryptophan Hydroxylase, the 5-HT synthesis rate-limiting enzyme) in mutant and non-mutant CF and SF at 5mpf. Again, morph-specific and region-specific differences were observed. TPH activity was strongly decreased in mutant CF (Fig 4). Of note, the reduction was more important in the anterior part (−53%) than in the posterior part (−28%) of the brain. In surface fish however, the reduction of TPH activity was observed only in the anterior part of the brain, and was more modest (−15%). Importantly, this result is fully consistent with the fact that 5-HT levels in the posterior brain of cavefish are unaffected by the P106L *mao* mutation, conversely to surface fish. Finally, TPH activity was identical in non-mutant SF and CF. Thus, we concluded that a significant compensation of the effects of the mutation exists at the 5-HT synthesis level, that this compensation is stronger in cavefish, and that the anterior and posterior 5-HT nuclei behave differently in this respect.

**Figure 4.**
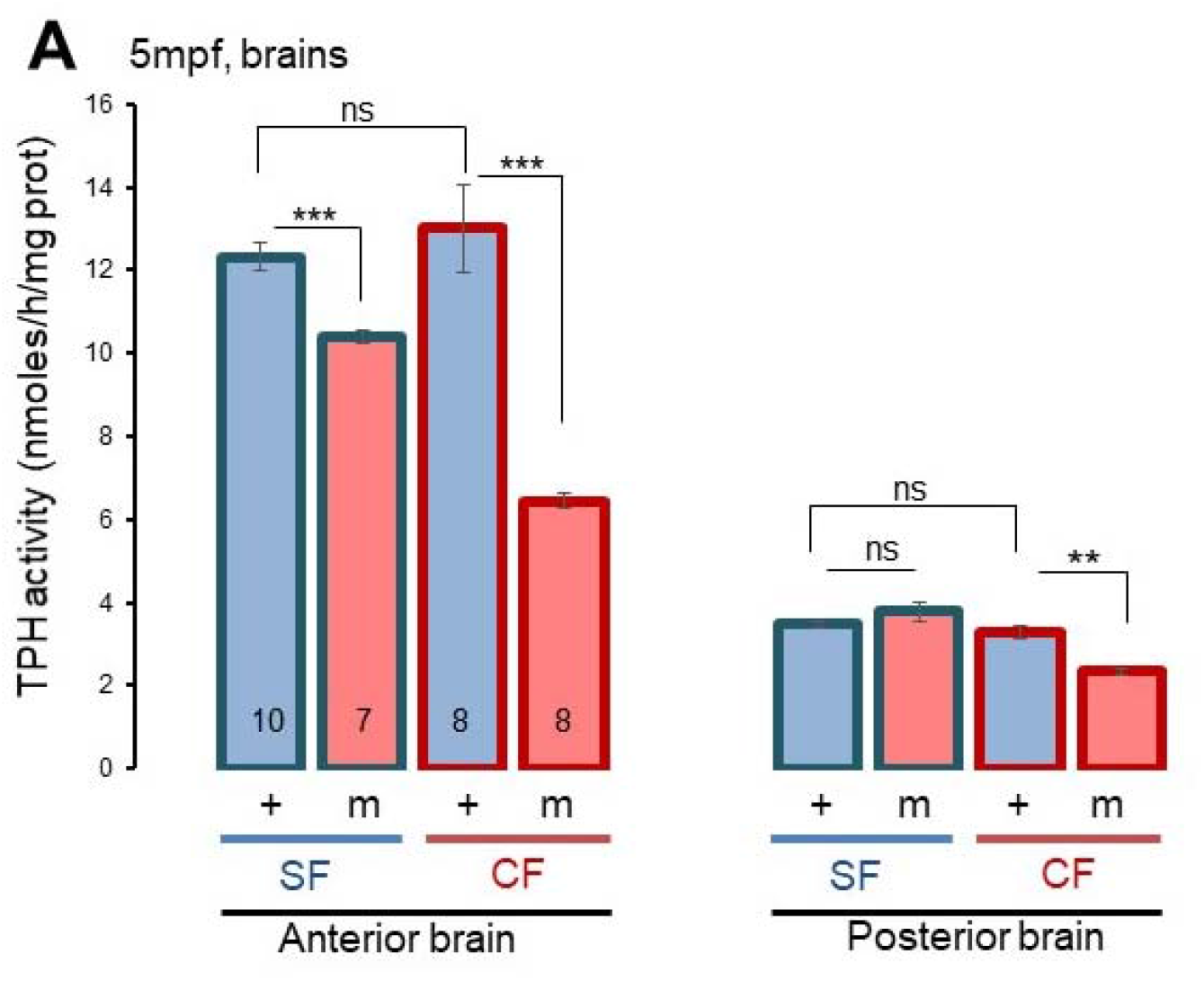
Tryptophan hydroxylase activity in *mao* P106L mutant or non-mutant cavefish and surface fish. (A) TPH activity was measured in the anterior and posterior brain of 5 month-old individuals, SF and CF, mutant (m) and non-mutant (+).

#### A chronic treatment with Deprenyl does not mimic the P106L genetic mutation

We next asked to what extend the effects of the P106L mutation could be compared to MAO inhibition by drug treatment. We thus analyzed the effects of Deprenyl, a selective and irreversible MAO inhibitor, on brain neurochemistry.

Acute treatment on 5 months-old adult cavefish increased 5-HT levels by 2.03X and 2.36X in non-mutant CF, and by 1.35X and 2.47X in mutant CF, in the anterior and the posterior part of the brain, respectively (Fig 5). As expected, the treatment also decreased 5HIAA levels. The modifications caused by acute MAO inhibition by Deprenyl, at the concentration used (10μM), were stronger than those observed with the P106L mutation. Of note, this acute pharmacological treatment also increased 5-HT levels in the posterior part of the brain, contrarily to the mutation (Fig 5). On the other hand and surprisingly, Deprenyl had no effect on NA and DA levels, but their metabolites were systematically and significantly decreased. Taken together, these results showed that an acute MAO inhibition had markedly different effects on brain neurochemistry than a chronic MAO deficiency due to a partial loss-of-function genetic mutation.

**Figure 5.**
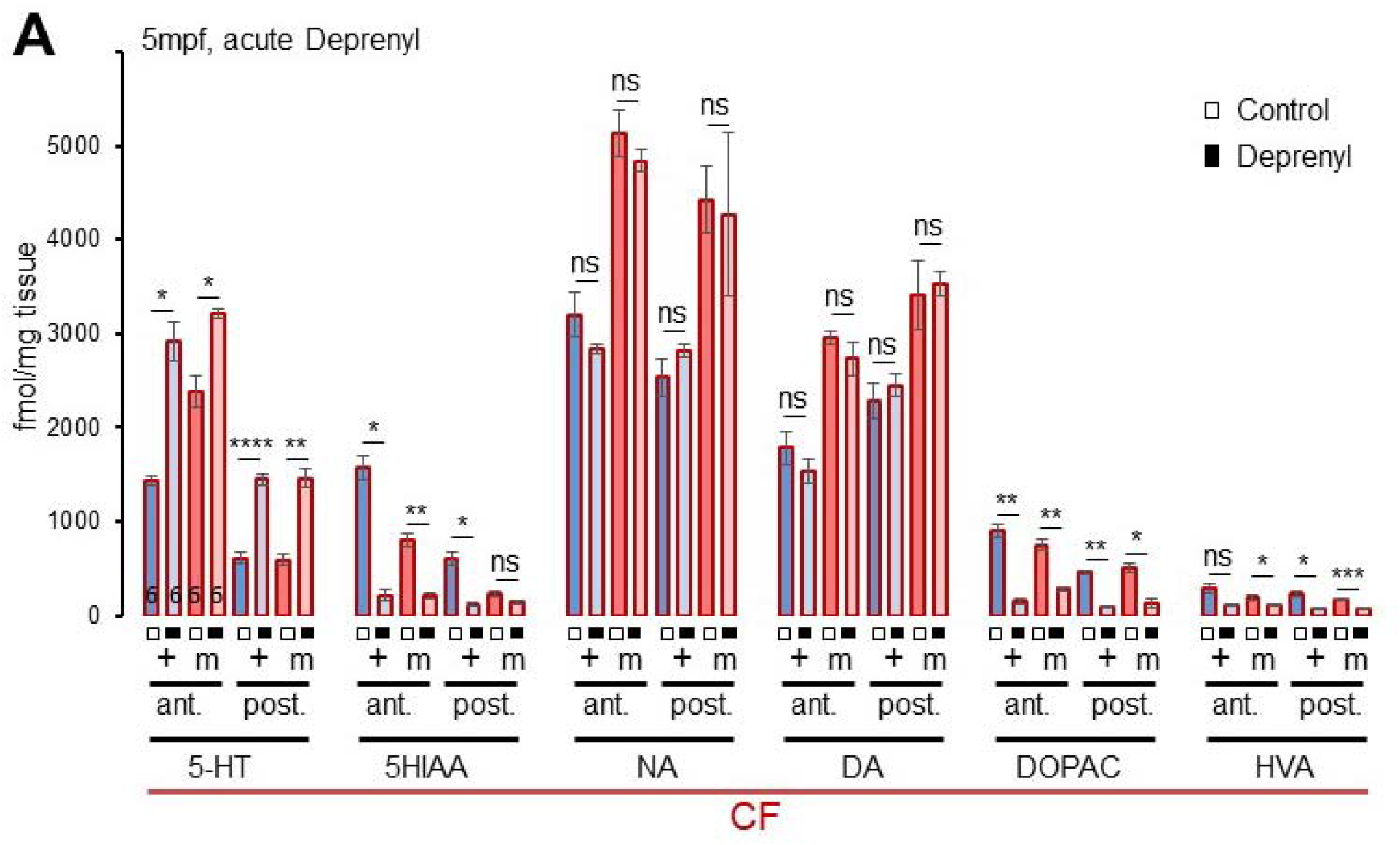
Effects of acute Deprenyl treatment on cavefish brain neurochemistry. (A) Monoamine and metabolites levels in the anterior and posterior parts of the brain of 5 month-old fishes, after acute 10μM Deprenyl treatment (black squares) or not (white squares); n=6 each. Fish used are CF, both mutant (m) and non-mutant (+).

## Discussion

Our data provide insights, 1) for understanding *A. mexicanus* cavefish phenotype and evolution, 2) for understanding the specificities of fish monoaminergic and particularly serotonergic systems, and 3) for understanding MAO-dependent regulation of brain neurochemistry in general. These 3 aspects are discussed below (Fig. 6).

**Figure 6.**
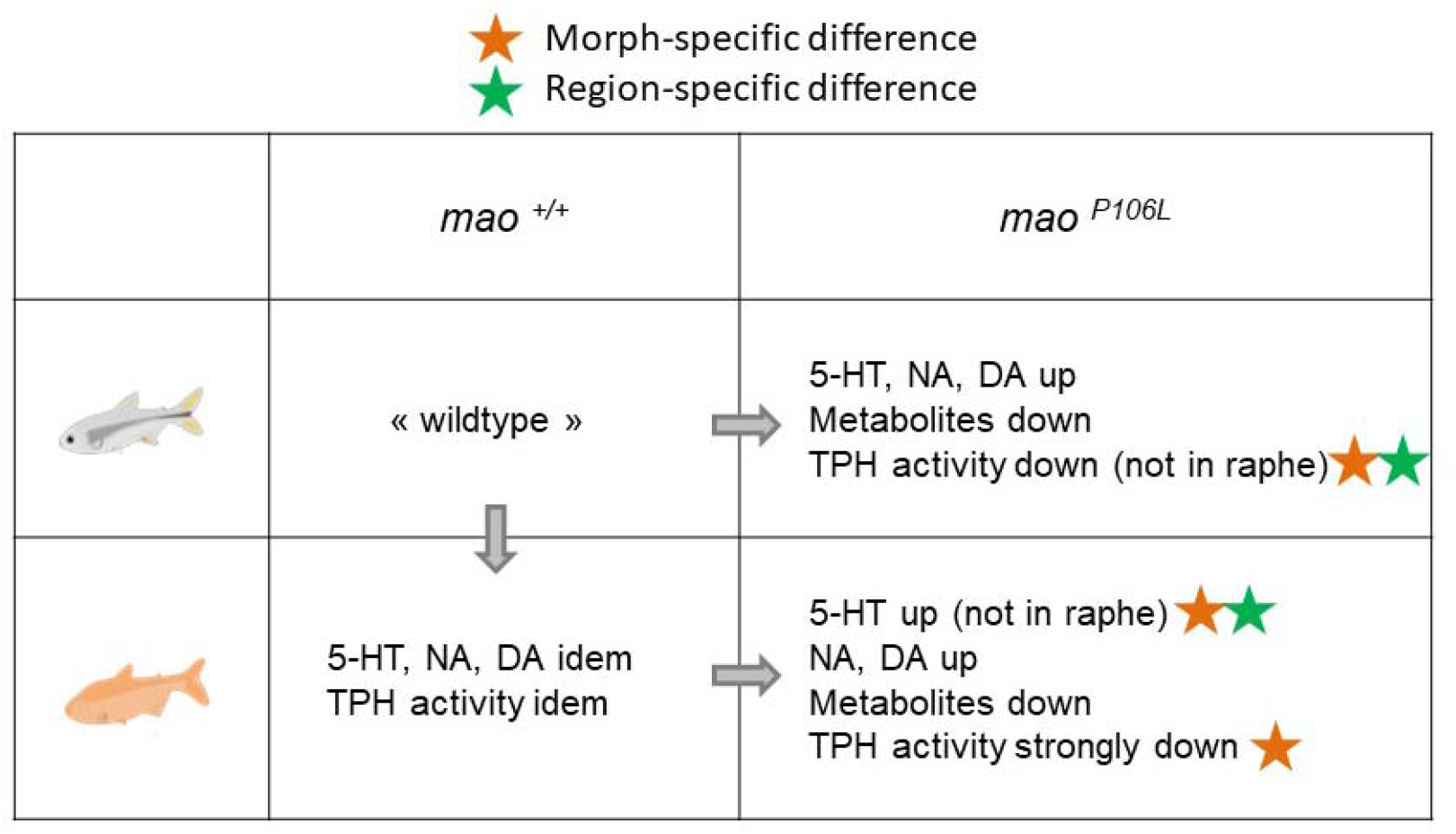
Summary of the results. The variations in brain neurochemistry observed between surface fish and cavefish on one hand, and non-mutants and mutants on the other hand, are summarized. The four-ways comparisons allows inferring genotype-specific, morph-specific, and brain region-specific differences.

### *mao* P106L: specific considerations related to cavefish biology

The first crystal structures of human MAO-B and rat MAO-A were obtained in 2002 and 2004, respectively (Binda *et al.* 2002; Ma *et al.* 2004), and subsequent, increasingly higher resolution structures (De Colibus *et al.* 2005; Son *et al.* 2008) helped the development of effective and selective inhibitors. The C-terminal region of the protein corresponds to transmembrane alpha-helices that anchor the enzyme to the mitochondrial outer membrane. The active site is a flat hydrophobic cavity occupied by the covalently-bound FAD coenzyme at the distal end. To access the active site, the substrate (or inhibitor) must negotiate a turn near the intersection of the enzyme with the surface of the membrane, and enter through 3 loops which form a narrow path. In the human MAO-A, the entrance is surrounded by residues V93-E95, Y109-P112, and F208-N2012 which lie in 3 different loops, and therefore loop flexibility is critical for opening the entry for substrates (Son *et al.* 2008). In the *A. mexicanus* MAO, we have found that the overall 3D structure of the enzyme is very well conserved. However, the P106L mutation is precisely located on one of these 3 entrance loops, and confers additional rigidity to the loop, as suggested by predicted extra beta-strands on the secondary structure. Therefore, we propose that the P106L mutation in cavefish MAO could impair the conformational fluctuations necessary for loop flexibility and substrate entrance into the enzyme active site. This hypothesis derived from structural *in silico* modeling is well supported by computational predictions on the pathogenicity of the mutation obtained from the machine-learning based predictor MutPred2 (Pejaver *et al.* 2017) and, most importantly, by biochemical evidence showing that *A. mexicanus* MAO enzymatic activity is reduced by two thirds by the P106L mutation. Thus, P106L corresponds to a partial loss-of-function mutation, and the oxidative deamination of monoamine neurotransmitters and biogenic amines must be strongly impaired in cavefish metabolism.

The cave-dwelling morphs of *A. mexicanus* are albino. Their lack of melanin pigmentation is due to a mutation in the gene *Oca2* (*Ocular and cutaneous albinism 2*; involved in the transport of melanin precursor into melanosomes)(Protas *et al.* 2006). It was suggested that a potential benefit of albinism in cavefish could be to provide a surplus of L-tyrosine, the common precursor of the catecholamine and melanin synthesis pathways, and that this could explain the higher levels of DA and NA in cavefish brains (Bilandžija *et al.* 2013). In support of this hypothesis, morpholino knock-down of *Oca2* expression in surface fish embryos decreased their pigmentation as expected, and simultaneously increased the L-tyrosine and DA content in 3 days-old larvae (Bilandžija *et al.* 2013). In contrast to this hypothesis though, here we bring strong, genetically-based evidence at different stages, from 6 days to 1 month and up to 5 months of age, that the differences in brain 5-HT and DA content between SF and CF is fully and solely due to the P106L *mao* mutation. Indeed, monoamines and metabolites HPLC patterns and levels were similar in SF and non-mutant CF, and altered in mutant CF, thus pointing to the causal role of *mao* P106L. Of note, the slightly increased DA levels in non-mutant CF as compared to SF (+9%) we observed at 6dpf may correspond to a transient and limited contribution of the *Oca2* mutation to increasing DA levels in cavefish at early developmental stages, when monoaminergic systems are already developed but probably not fully mature (Elipot *et al.* 2014; Pierre et al 2020). For comparison, even at this early larval stage, the *mao* mutation alone increases DA levels by 30% (Fig. 3A). And at later stages, the *mao* P106L contributes to 100% of the observed differences, and for all monoamines.

In a companion paper on the neuro-behavioral effects of the *mao* P106L mutation, we have found surprisingly little effect on the cavefish phenotype, besides a clear involvement of altered brain monoamine homeostasis in the “stressability”, or amplitude of response to a stress represented by a change of environment (Pierre et al., 2020). This is at odds with the major cognitive and behavioral deficits caused by the only two *MAO-A* mutations (Glu313Stop; Cys266Phe) discovered so far in two human families (Brunner *et al.* 1993; Piton *et al.* 2014), and with the two-third reduction of MAO enzymatic activity and the ~33% increase in monoamine brain levels we have observed in mutant fish. This discrepancy raises two points of discussion. First, there must be multiple and complex compensatory plastic mechanisms occurring in the brain and body of cavefish, and which explain how they can cope with their *mao* genetic mutation. It will be important to study these plastic changes, as they can give valuable information on general mechanisms of homeostasis. Here, we have found that this includes, at least in part, a compensation at the level of neurotransmitter synthesis, as shown by reduction of TPH activity in P106L *mao* mutants. Second, it is well known that cavefish display metabolic alterations, accumulate fat, and suffer hyperglycemia and diabetes due to a mutation in insulin receptor (Aspiras *et al.* 2015; Riddle *et al.* 2018; Salin *et al.* 2010). As a potential role of MAO in metabolic and cardiovascular diseases has recently been raised (Deshwal *et al.* 2017; Shih 2018), it will be important as well to examine a possible contribution of *mao* P106L to non-neural, metabolic phenotypes of cavefish.

### General considerations on MAO and 5-HT system in fish

There are two major differences between mammals and fish concerning MAO and the 5-HT system (Lillesaar 2011). Fish possess only one MAO as opposed to MAO-A and -B in mammals, and they possess clusters of 5-HT neurons both in the hypothalamus and in the raphe nucleus, as opposed to raphe only in mammals. *Astyanax mexicanus* does not escape the rule in these respects (Elipot *et al.* 2013; Elipot *et al.* 2014; Pierre et al., 2020).

Fish MAO has been suggested to have properties closer to mammalian MAO-A, including a suspected poor affinity for dopamine as a substrate. This is because inhibition of MAO by deprenyl in larval zebrafish strongly elevates 5-HT levels, but not DA or NA levels (Sallinen *et al.* 2009). The same holds true in *A. mexicanus* (Fig. 5). However, in the present study after acute deprenyl treatment, we found no change in DA and NA levels like in zerafish, but a very strong decrease in the levels of DA metabolites DOPAC and HVA, which is counterintuitive if DA is not a MAO substrate (NB: DOPAC can only be produced by MAO-dependent degradation, therefore its presence suggest that DA is degraded by MAO). This discrepancy between the variations of metabolites and neurotransmitters cannot be explained easily either if one considers the alternative catabolic pathway for DA, through COMT (catechol-O-methyltransferase). Even if the fish MAO was to degrade efficiently dopamine, a conundrum would remain. Why and how do DA and NA levels remain stable when their metabolites vary? An unknown, fish-specific mechanism must exist, which buffers the expected rise in DA and NA when MAO is pharmacologically inhibited – but not when it is genetically mutated. Indeed in the case of the P106L cavefish mutation, neurotransmitters and their metabolites show logical, inversely correlated variations, which also tends to suggest that DA and NA are *bona fide* MAO substrates in fish. These observations suggest that a “reference value” for the level of these catecholamines might exist, and would depend on the *mao* genotype.

Interestingly, in our experiments the hypothalamic and the raphe serotonergic neurons did not respond in the same manner to MAO perturbations (Fig. 6). In fact, while hypothalamic 5-HT and 5HIAA levels varied according to the *mao* genotype for the two morphotypes (CF and SF) and the two genotypes (mutant and non-mutant), in the posterior brain the 5-HT levels remained the same while the 5HIAA levels were lower specifically in *mao* P106L CF mutants (Fig. 3C). This confirmed and reinforced previous results (Elipot *et al.* 2014), and suggested a compensation of the effects of the *mao* P106L mutation, in the CF raphe, to tightly regulate the 5HT levels. The decreased TPH activity we have observed in the posterior brain of the mutant cavefish -but not the mutant surface fish-may participate to the process. Furthermore, in CF an acute MAO inhibition by deprenyl treatment did modify 5-HT raphe levels whereas a chronic inhibition by the P106L genetic mutation did not. This suggests that the compensation in the raphe, if it exists, takes time to set up.

During embryonic development in zebrafish, the specification of the 5-HT neurons of the hypothalamus and the raphe is controlled by different transcription factors: pet1 in the raphe (Lillesaar *et al.* 2007) like in mammals, *versus* Etv5b in the hypothalamus (Bosco *et al.* 2013), suggesting that the serotonergic identity can be acquired by convergent mechanisms. Moreover, hypothalamic and raphe 5-HT neurons express distinct paralogs of the 5-HT pathway markers: *tph1* and *sertb* in the hypothalamus, *tph2* and *serta* in the raphe (Lillesaar 2011). The timing of their differentiation is also different. We propose that the different ways the anterior/hypothalamic and the posterior/raphe neurons respond to the partial loss-of-function *mao* mutation in cavefish must be related to their distinct serotonergic identity conferred by their distinct embryonic origins. Hence, the cavefish used as a model for evolutionary biology can bring interesting insights, in a comparative perspective, to basic knowledge on neurotransmission homeostasis.

### General considerations on MAO-dependent brain chemistry

In the cavefish brain, as expected, the P106L *mao* mutation increased the levels of 5-HT, DA and NA (except for 5-HT in the raphe, discussed above and below) and decreased the levels of their metabolites. These results are consistent with those found in MAO-A simple mutant or MAO-A/B double mutant mice, where the NA, 5-HT and DA levels are increased, and the 5-HIAA and DOPAC levels are decreased (Cases *et al.* 1995; Popova *et al.* 2001; Chen *et al.* 2004).

Here, we would like to discuss the sometimes-contrasting results observed in P106L *mao* mutants *versus* acute deprenyl treatment. The P106L *mao* mutation changed DA and NA levels whereas acute deprenyl treatment did not. The later is consistent with Sallinen *et al.* (2009) who found an increase of 5-HT levels but not NA and DA levels after deprenyl treatment in larval zebrafish. Therefore, pharmacological and genetic interference on MAO function does not have the same outcomes on fish brain neurochemistry. In mammals too, such contrasting effects have been observed. Let’s take the raphe as an example to discuss the possible underlying mechanisms.

The mammalian raphe receives noradrenergic projections from the locus coeruleus (Levitt and Moore 1979; Baraban and Aghajanian 1981) and is under tonic activation via α1-adrenoreceptors (Baraban and Aghajanian 1980). On the other hand 5-HT1A autoreceptors and α2-adrenoreceptors mediate an inhibitory effect on raphe 5-HT neuronal activity (Andrade *et al.* 2015; Tao and Hjorth 1992; Numazawa *et al.* 1995; Raiteri *et al.* 1990; Trendelenburg *et al.* 1994; Starke *et al.* 1989). The 5-HT2A/2B/2C, 4 and 6 receptors (Belmer *et al.* 2018; Lucas and Debonnel 2002; Liu *et al.* 2000; Boothman *et al.* 2006; Brouard *et al.* 2015), the endocanabinoid system (Haj-Dahmane and Shen 2011), and GABAergic neurons (Hernández-Vázquez *et al.* 2019) also modulate the 5-HT system. Little is known about serotonergic modulation in fish, but the teleost and mammalian monoaminergic systems share numerous similarities (Panula *et al.* 2006) and their raphe 5-HT neurons can be considered homologous. Among others, projections to the raphe from the locus coeruleus (Ma 1994), expression of α2-adrenoreceptors and 5-HT1A receptor in the raphe (Ampatzis *et al.* 2008; Norton *et al.* 2008) have been described in fish.

In mammals, acute MAO-A inhibition causes a decrease of firing activity of 5-HT dorsal raphe neurons (Blier and De Montigny 1985). The underlying suggested mechanism is that MAO-A inhibition would increase levels of synaptic 5-HT, which normally activates inhibitory 5-HT1A autoreceptors (Haddjeri *et al.* 1998). Along this line, 5-HT would accumulate both because of the reduction of 5-HT degradation by MAO-A after re-uptake, and because of the reduction of 5-HT released and re-uptaked due to the decrease of firing activity (Haddjeri *et al.* 1998).

Conversely, with a chronic MAO-A inhibition (chronic pharmacological treatment or MAO-A knockout), the increase in 5-HT levels are more moderate (Haddjeri *et al.* 1998). Also, a long-term treatment with befloxatone (MAO-A inhibitor) induces a desensitization of inhibitory α2-adrenoreceptors (Blier and Bouchard 1994; Mongeau *et al.* 1994; Owesson *et al.* 2002), and decreases their density (Cohen *et al.* 1982). Moreover, several studies suggested that a chronic MAO inhibition decreases the inhibition of firing rate mediated by 5-HT1A autoreceptors (Owesson *et al.* 2002; Palfreyman *et al.* 1986; Evrard *et al.* 2002). Together these effects lead to suppression of 5-HT neurons inhibition. Some studies reported a complete recovery of the raphe firing activity with chronic treatment by clorgyline (MAO-A inhibitor) (Blier and De Montigny 1985), while others reported that the firing activity of 5-HT neurons was still lowered by 40% in MAO-A KO mice (Evrard *et al.* 2002). In sum, different responses of the 5-HT raphe system to acute or chronic MAO inhibition are described in mammals: decrease of 5-HT raphe activity with acute inhibition, and complete or partial re-establishment of 5-HT raphe activity with chronic inhibition. Equivalent studies were not performed in fish, but they give possible explanations to the different effects of acute (deprenyl) and chronic (P106L mutation) MAO inhibition on 5-HT raphe levels in fish.

Finally, we discovered a compensation to the effects of the P106L mutation by a decrease of tryptophan hydroxylase (TPH) activity in mutant CF, and in both parts of the brain. This could attenuate the increase of 5-HT levels in mutant CF, and explain why MAO enzymatic activity is proportionately more reduced than monoamine levels are enhanced in mutant brains. Of note, we did not detect such a significant reduction of TPH activity in our previous study when we used cavefish samples that were probably not genetically homogeneous (Elipot et al., 2014). This highlights the importance of re-examining all parameters in genetic lines with known genotypes, and the potential hidden pitfalls of working on natural, polymorphic populations. Further studies are needed to know if the expression of *tph* is decreased and, more generally, whether and how the brain transcriptome as a whole is profoundly affected or dys-regulated in the *mao* mutant context. The mechanism of the “TPH compensation” is also unknown, but one possibility could be that the excess of synaptic 5-HT leads to a negative feedback on 5-HT synthesis. For example, increased TPH activity has been shown after destruction of 5-HT-containing nerve terminals in mice (Stachowiak *et al.* 1986). But conversely, Popova *et al.* (2001) showed that in the MOA-A KO mice, the TPH activity in frontal cortex, hippocampus and amygdala was increased, which did not attenuate but rather aggravated the effects of MAO-A deficit.

Future experiments will have to test whether the synthesis of DA and NA is modulated as well by the P106L mutation, which could compensate the increase of DA and NA levels in *mao* mutants. Indeed, in mammals, several studies showed modifications of TH activity after acute or chronic inhibition of MAO-B (Vrana *et al.* 1992; Lamensdorf and Finberg 1997).

## Conclusion and perspectives

Using the cavefish as a natural mutant, we have uncovered several novel aspects of monoaminergic regulation in fish. Fish are doubly special as compared to mammals because they have only one *mao* gene but possess diversified and diffuse clusters of monoaminergic neurons in their brains. However, they can serve as valuable models because most characteristics and most functions are shared with mammals. Our findings open a width of perspectives for future studies. This will include the understanding of the enigmatic differences in homeostatic properties between hypothalamic and raphe serotonergic neurons, or the deciphering of the probably complex plastic compensatory mechanisms that occur in the cavefish brains and allow them to thrive in caves despite a deleterious mutation in a major enzyme, MAO.

